# Predicting Speech in Noise Perception using Measures of Neural Entrainment

**DOI:** 10.64898/2026.09.08.750184

**Authors:** B.M. Bormann, Z. Balky, C. Li, A. Ghasemzadeh, Kelsey Mankel, Daniel C. Comstock, S. Das, R.S. Whittle, H. Brodie, D. Sagiv, L.M. Miller

## Abstract

Speech perception ability in a cocktail party environment is highly variable, even for people with clinically healthy hearing. The mechanisms of these differences are poorly understood. A possible reason for these differences could be due to the neural mechanisms of auditory attention. This investigation aims to quantify how well a person is able to neurally entrain to a continuous narrative using a temporal response function (TRF), then evaluate if that TRF measure is correlated with an individual’s performance on a selective attention and narrative comprehension task. This was accomplished using a cohort of twenty-five young and healthy listeners who completed speech-in-noise (SIN) perception tasks while their neural activity was recorded with EEG. This EEG data and regressors (speech envelope and gammatone spectrogram) of an auditory stimulus was used to train TRF models. TRF cross validation correlation coefficients were then used in multiple regression models to predict a participant’s SIN performance. Task performance had a significant positive relationship with the neural entrainment to an attended stimulus and a negative relationship with the neural entrainment to an unattended stimulus. This was observed with both the selective attention and narrative comprehension measures. The difference of the attended and unattended correlation coefficients were also significantly related to increased task performance. These measures could be a potential biomarker to predict an individual’s speech perception in a noisy environment.

## Introduction

The ability to perceive speech is highly variable across individuals, even those with clinically normal hearing.^1,2^ This variability is even more pronounced in noisy environments where a person must attend to a target while ignoring competing stimuli.^3^ The reason for this difference of perceptual ability is unclear. It is possible that this variability is due to how different people utilize an array of cognitive and neural resources to robustly represent a target stimulus in their brain while simultaneously suppressing the representation of the masking stimuli. This investigation aims to quantify how well a person’s brain is able to entrain to an external auditory stimulus using a temporal response function (TRF) and then see if that measurement of neural entrainment is related to how well a person is able to perceive speech in a cocktail party environment.

### Temporal Response Functions and Auditory Attention

Auditory attention has been extremely well studied using a wide range of electroencephalography (EEG) methods. The most prevalent of those methods is event related potential (ERP) analysis because it is an efficient way to isolate extremely small changes in neural activity due to attention.^4^ A drawback of this method is that it is dependent on averaging the neural activity across many short and repetitive trials. This is a fantastic tool when investigating mechanisms that change over the factor of milliseconds but lacks the ability to investigate the fluctuations for a process that last for minutes or longer. This is why TRFs have been an extremely powerful tool for auditory neuroscientists investigating how the brain responds to more naturalistic speech stimuli over time.

TRF is a linear fit model that can be trained on the relationship between a stimulus and a response over time.^5,6^ This model can then be used to predict either the neural response to a specific stimulus or to predict what stimulus would occur for a given response. The majority of the literature in this field has focused on using forward TRF models. This method predicts the neural response of an EEG waveform based on the stimulus features of an external auditory stimulus. This is an effective method to capture how modeled brain activity will fluctuate across the whole scalp to better understand how the brain represents different auditory features. These predicted modulations of the EEG waveforms are then cross-referenced across the actual EEG recordings to determine how well correlated the two are. This can be used to show how well the brain represents the different aspects of the stimulus. A classic example of this is how the brain can entrain on the audio envelope of a naturalistic voice.^5,7^ The strength of these correlations between the predicted and actual EEG activity can vary based on high-level cognitive processes, like attention.^8^ In a study looking at the correlation between the predicted EEG activity and the actual EEG activity for both attended and unattended speech, it was observed that the correlation between the predicted and actual EEG for the attended stimulus was significantly greater than the correlation for the unattended stimulus.^9^ This indicates that an attended stimulus will be represented in the brain in a way that is more easily decoded by the TRF model. This most likely means the brain is better entrained on the attended stimulus than the unattended stimulus. This relationship is consistent enough that it can be used to determine if a stimulus is attended to or ignored by a participant using only the TRF cross validation correlation values.^10,11^

Forward TRF models have also been used to investigate how the brain represents an auditory target stimulus in different noisy environments. It has been demonstrated that the TRF filter morphology is related to the number of words a person can understand in a noisy environment.^12^ It has also been observed that success in perceiving words in short sentences presented with different degrees of noise is correlated with the neural entrainment to the speech envelope, that relationship of neural entrainment and successful listening is further influenced by age.^13^ This shows that the intensity of competing stimulus influences how the brain can entrain on an external stimulus. Investigations have also shown that people with impaired speech perception show differences in their cortical tracking of speech in noisy environments.^14^

While TRF has been shown as an effective method to investigate auditory attention to continuous speech, there is very little known about how these models can be used to characterize a person’s ability to selectively attend to speech and to comprehend the information being communicated. This manuscript aims to fill that missing link. To accomplish this, we aim to use the correlation coefficient of a backward TRF model to quantify how well a person can entrain to a target stimulus and how well they suppress their entrainment to a masking stimulus. We then aim to evaluate if the strength of those correlations is related to a person’s performance on a selective attention task and narrative comprehension task.

### Predicting Speech Perception with Neural Entrainment

We posit that a person’s performance on a selective attention task and narrative comprehension task will have a positive relationship with the attended condition correlation coefficient while having a negative relationship with the unattended condition correlation coefficient. Furthermore, the difference of the attended and unattended correlations will also have a positive relationship with a person’s performance on that task. Implying that increased task performance is not just dependent on the degree of entrainment for the attended condition, but the individual’s ability to differentiate the neural representation of separate auditory signals.

The reason we selected using the backwards model, rather than a forwards model, is because we are aiming to find an informative and accessible biomarker that captures the variance of the whole brain. The backwards model reverse engineers the features of a stimulus based on the neural activity of all the electrodes, thus providing a holistic representation of the brain in one value. We believe this will be a practical measurement for future researchers and clinicians to better understand how people are able to perceive speech in a noisy environment. We hope to determine if a backwards TRF correlation coefficient can be used as a biomarker to categorize a person’s ability to perceive and comprehend speech in a cocktail party environment. This measure could facilitate a better understanding of the neural mechanisms responsible of the interindividual variability of speech perception. In addition, a possible biomarker could also be used to better conceptualize the neural activity of people with different perception deficits like attention-deficit hyperactivity disorder (ADHD), central auditory processing disorder (CAPD), and hidden hearing loss.

## Methods

### Participants

This analysis was performed on the same collection of participants from the investigations performed by Shebahi et al. 2025 & Mankel et al 2026. Twenty-nine adults were recruited to participate in the study. Four of those subjects did not complete the study due to being ineligible or left for personal reasons. Further details about participant exclusion criteria can be found in Shehabi et al & Mankel et al.

The cohort analyzed was a collection of twenty-five adults (15 females & 10 males) who aged from 18-38 years old (M= 22.88 ± 4.98 SD). All participants had clinically normal hearing ability which was determined by having equal or less than 25 dB HL air conduction pure tone audiometric (PTA) thresholds averaged across 500, 1,000, and 2,000 Hz (M = 6.933 ± 3.8767 dB HL).

### Project Overview

This investigation is focused on a subdivision of data from a larger project aiming to better characterize the clinical, cognitive, & neural measures of listening ability to continuous speech. This larger project was a three-part study which included an hour long clinical audiological assessment, a 2-hour session where participants completed a battery of cognitive and psycho-acoustic tasks, and a 2.5-hour electroencephalography (EEG) session where they completed a continuous speech spatial attention task while their neural responses were recorded. Participants could complete these sessions over the span of several days. Participants who decided to complete multiple sessions in the same day were encouraged to take an approximately 1-hour break to feel prepared for the next session.

The data set being used in this project was only collected during the clinical assessment and the EEG session. PTAs for the participants were collected during the audiological assessment to screen out participants with different forms of hearing impairments and to characterize their clinical hearing ability. The rest of the data utilized for this analysis was collected during the EEG session. This includes the EEG activity used to train the temporal response functions (TRF) and the behavioral performance from the spatial auditory attention task and narrative comprehension task.

### Chirped Speech (Cheech) Stimulus

To investigate the neural activity linked to processing continuous speech across the ascending auditory pathway, a specially designed vocal stimulus was presented to the participants. Chirped speech (Cheech) is a method in which the glottal pulses of speech are replaced with the CE-Chirp which are extremely fast frequency sweeps from 200-10,000 Hz. This modification elicits a robust and consistent auditory brainstem response (ABR) while still allowing the recording of middle latency responses (MLR) & late latency responses (LLR) that would be elicited from typical speech stimulus. This stimulus enables us to assess the auditory pathway, from cochlea to cortex, using one ecologically relevant stimulus.

### Spatial Selective Auditory Attention Task

The stimulus used for this analysis is the same as Shehabi et al and Mankel et al, which was a Cheeched voice narrating 7.5-minute-long fairy tales. These fairy tales were edited to include 25 monosyllabic target color words added organically to the existing text (e.g. “Dear father, bring me a *pink* rose”). The voice speaking the narrative would appear to be either 15 degrees to the left or right side of their head using a head-related transform function (HRTF). The locations would then switch to the opposite side approximately every 6 seconds to mimic a dynamic listening environment and to maintain active engagement of the stimuli. Further information about the preparation of the Cheech stimulus can be found in Shehabi et al and Mankel et al.

Participants listened to the stimuli during either a speech-in-quiet (SIQ) condition where they only heard one narrative and no competing stimulus being presented, or a speech-in-noise (SIN) condition where they would attend to a narrative while another masker narrative was played concurrently. The narratives of the SIN conditions were presented with different gendered voices and spatially separated from each other to help reduce confusion. Furthermore, participants had a visual cue on the monitors in front of them to help them differentiate the voices. The side with the target narrative had a large white “+” on the center of the screen which was aligned with the HRTF of the voice. This meant the voice appeared to be coming from the “+” on the screen, while on the opposite side the masker voice has either a “<” or “>” on the screen which was pointing to the side of the target speaker. This was to help remind the participant to direct their attention to the side of the target narrative (aka the “+”). Stimuli presentation was controlled by a computer with a Linux operating system running an in-house MATLAB code using the Psychtoolbox package. The presentation computer relayed the audio information through a Hammerfall audio card, MRE Fireface UFZ interface, and RME ASI-2DAC FS amplifier. The amplifier would then play the audio over a set of ER-2 insert earphones that were shielded to prevent interfering with the EEG recordings.

Subjects participated in a continuous speech selective attention task while EEG was used to record their neural activity during the task. Participants sat in a sound dampening recording chamber with the lights turned off. Participants would be instructed to attend to a continuous narrative being played over a pair of in-ear headphones. They were informed that they needed to listen specifically for color words (e.g. “red”, “blue”, “pink”, etc.) which were naturally imbedded into the narrative. They would then press a button each time they heard one of the target color words while ignoring any color words in the masker narrative. Their accuracy and reaction time (RT) for each button pressed was recorded and interpreted as measures of their selective attention. Participants were also instructed to attend to the events of the narrative because they would be completing a series of multiple-choice questions about the narrative. These questions were used to quantify the subject’s narrative comprehension ability.

Participants listened to six different story conditions over the course of their EEG session where different factors such as the number of speakers, the gender of the speaker, and additional transformations to the Cheech stimulus were varied. This analysis is specifically analyzing data from two of those six conditions. The stimuli from these two blocks are identical to each other. They are the same SIN condition where the participant is attending to a target narrative on one side of their head, while simultaneously ignoring another masker narrative on the opposite side. Each block was comprised of two different fairy tales being narrated by a male Cheeched voice and a female Cheeched voice. While the stimuli were identical, the participants were instructed to attend to the male voice for one block, while being instructed to attend to the female voice during the other block. This allows us to isolate the effects of attention while holding other factors such as narrative and question difficulty constant across blocks. This analysis was not repeated for the female voice due to the significantly different performance across the selective attention and narrative comprehension tasks. It is unclear if behavioral performance differed due to the difference in speaker gender, the narrative itself, or the difficulty of the task. All analysis performed were with the male speaker to isolate the effects of attention while holding other variables constant.

### EEG- Data Collection

Participants neural activity was recorded using 64-channel cap electrodes in the International 10-20 standard layout. Eight additional electrodes on and around their ears were used for another ongoing project. These additional electrodes were placed on the earlobe, mastoid, posterior to the earlobe, and superior-anterior to the tragus near the zygomatic arch. All electrodes were connected to a BioSemi ActiveTwo amplifier with a sampling rate of 8192 Hz. EEG recording was performed on a PC desktop running Windows 10. Before recording, impedances for each electrode were reviewed and adjusted to ensure they were all below 20 µV when compared to the Common Mode Sense electrode. Impedance and EEG waveforms were reviewed using the ActiveView 2 software to quality assure the connection of each lead. Data was then collected and saved as an XDF file using open-source Lab Streaming Layer (LSL) software.

Brain Products StimTracks were used to time-lock the stimulus presentation and EEG recording by receiving trigger events from the separate target and distractor narrative. There the triggers were then relayed as trigger pulses to a TriggerBox (Brain Products GmbH) that communicated with the EEG amplifier and the data acquisition computer running LSL.

### EEG- Preprocessing

The XDF files with the EEG data were processed using a combination of EEGlab and custom code in MATLAB. Full details on EEG processing can be found in Shehabi et al, but the most important details for this report included the data being divided based on their respective conditions with a 10-second buffer before and after the end of the recording period. Those pruned stories then underwent a noncausal second-order 0.1 Hz Butterworth filter and were then cleaned of 60 Hz line noise using the cleanline plugin when needed. Independent component analysis (ICA) was used to remove possible artifacts such as eye blinks and heart beats. Noisy channels were then identified with manual inspection and then removed. This was then followed by a 1-50 Hz eight-order Butterworth bandpass filter in EEGlab. The EEG data was then downsampled to 64 Hz following ICA and filtered again using a 0.5-40 Hz eighth-order Butterworth bandpass filter. All electrodes were re-referenced to the average between the left and right ear lobe electrodes. Finally, for TRF analysis, EEG data was low-pass filtered to 8Hz using a zero-phase FIR filter, and then downsampled to 64Hz.

### TRF Model Training

We employed the multivariate Temporal Response Function (mTRF) implemented via the mTRFpy python package to calculate mTRF weight vectors for our backwards models. We calculated the audio envelope and 32 filter gammatone spectrogram as features of the audio stimulus, then we downsampled each feature to match the sampling rate of the EEG. These are the features regressed which were on, and then all 64 EEG channels are used to train a TRF model for each regressor. The mTRFpy package implements L2 regularization as a smoothing factor to prevent overfitting of the model [CITE]. We trained separate mTRF models for each participant during both attended conditions as well as unattended conditions. We swept window lags ([0,400], [10,410], [20,420], …, [100,500]), and regularization parameters (0.1, 1, 10, 100, 1000, 10000) in order to find values that led to the best cross-validated Pearson’s cross correlation on an individual participant’s data. For training, we split each story into 10 equal length segments, using 9 segments for training and 1 segment for testing with 10-fold cross-validation.

### Measures of Neural Entrainment

We used Pearson’s cross correlation (R-value) to determine how the predicted stimulus feature from the TRF correlated with the ground truth stimulus presented to the participant. This was calculated for the speech envelope and the gammatone spectrogram of the Cheeched audio. R-values from only the attended/unattended male narrative during the SIN conditions were used. The only difference across blocks is that the participant was instructed to attend to the male voice in one block while ignoring the female speaker, then performing the inverse for the other condition. This allowed us to isolate the neural difference due to attention because the auditory stimulus across the attended and unattended conditions are identical. The difference of R-values was then calculated (R-value for the attended condition minus the R-values for the unattended condition) to see if there is a relationship between the degree of neural entrainment across conditions with performance on a behavioral task.

### Outlier Assessment, Removal, & Replacement

Statistical analysis was performed using R version 4.5.2 (Posit Software, PBC) on a computer running MacOS (Tahoe 26.2). All data underwent an outlier analysis where all values 3 standard deviations from the mean were identified. Any of those outliers were then assessed for influence using a Cook’s distance analysis. Any values greater then 4 divided by the number of observations were removed. Any missing values due to technical issues or outlier removal were replaced using the multivariant imputation by chained equations (MICE) R package.

### Predictive Modeling of Performance

Mixed effects modeling was used to investigate the relationship between neural entrainment and performance on a spatial auditory attention task. These models were then used to predict a participant’s performance on the selective attention and narrative comprehension tasks based on R-values produced by the TRF. Linear mixed-effects regression was use for the color word hit reaction time because the measure was unbounded. The participant’s accuracy on the color word hit task and narrative comprehension task were both bounded measures (0-1) and several participants performed extremely well on the task causing possible ceiling effects. To minimize these ceiling effects, beta regression modeling was use for these behavioral measurements.

Two sets of regression models were performed. The first set of models were used to investigate the relationship of each of the behavioral measurements (hit reaction time, hit accuracy, and comprehension accuracy) with the backward TRF R-values for both the attended and unattended male SIN conditions in the same model. While the other set of models were used to probe the relationship of each behavioral performance metric with the difference of the attended and unattended R-values. Each of these models included the TRF predictor variables as well as the participant’s age and PTA to control for possible confounds.

Each of these models were used to predict a person’s behavioral performance on the spatial selective attention auditory task based on their backward TRF R-values. The predicted trends were then tested for significance using marginal means. These models were then plotted using the GGplot2 package in R with formatting performed in the BioRender web application.

## Results

### Task Performance

Participants completed a task where they attended to a spoken narrative on one side of their head while ignoring a concurrent masking narrative on the opposite side of their head. Subjects hit a button each time they heard a color word (e.g. “blue”, “pink”, etc.) naturally embedded into the target narrative. The accuracy and reaction time (RT) of these color word hits were used to quantify the person’s ability to selectively attend to a narrative while ignoring a distractor narrative. After listening to the narrative, they then answered a series of multiple-choice questions about the events that happened during the narrative. Performance on these questions were used to evaluate the participant’s narrative comprehension ability. Participants performed very well on all three measures. Their color word identification accuracy was 86.48 ± 10.16% (Figure 1.a) and their RT was 0.9232 ± 0.1605 seconds (Figure 1.b). Participants also scored 79.2 ± 22.72% (Figure 1.c) on the narrative comprehension questions.

**Figure 1.**
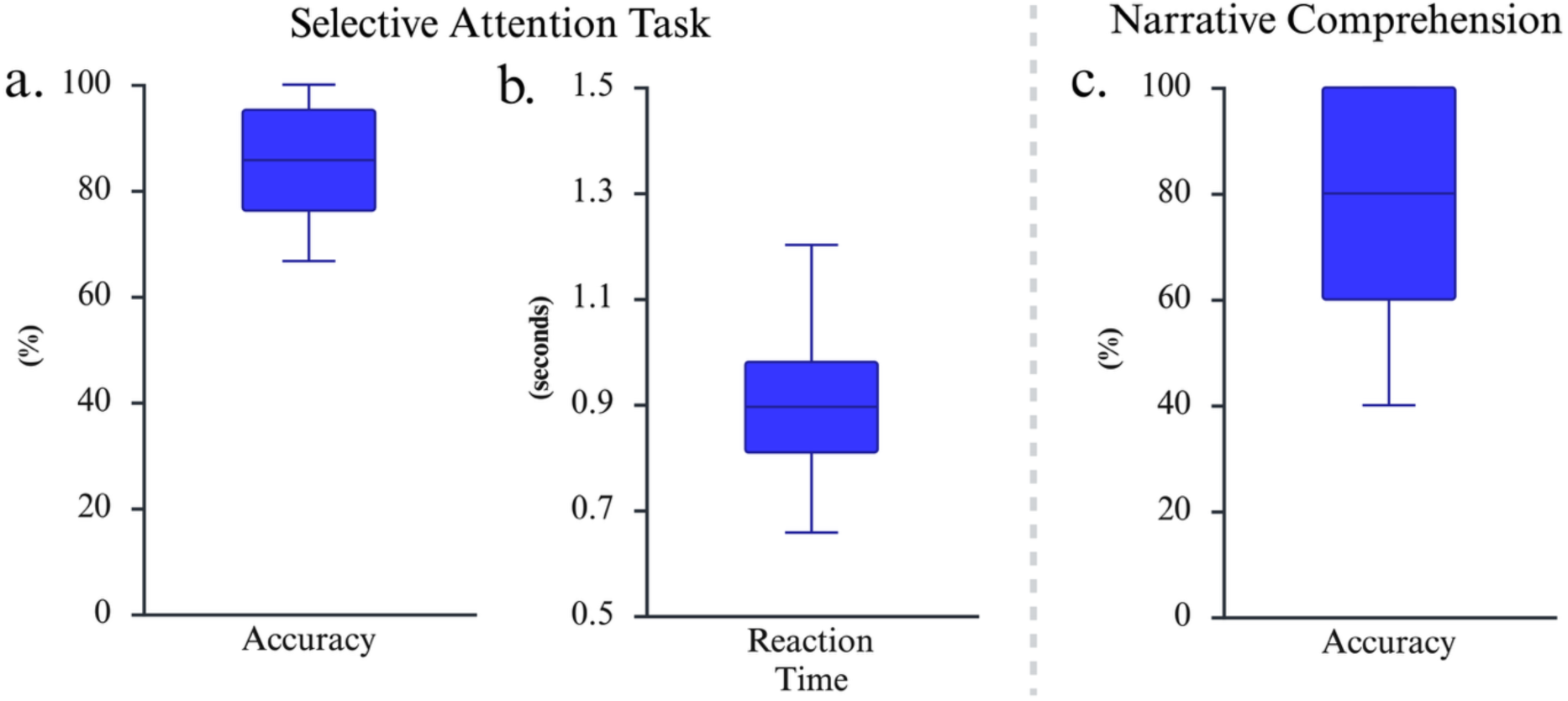
Box plots showing the performance of different speech-in-noise (SIN) tasks. Plots a & b show the participants’ hit accuracy and reaction time to a spatial selective auditory attention task and plot c show accuracy on a narrative comprehension task.

### TRF Measures of Neural Entrainment

Electroencephalography (EEG) was used to record the participant’s neural activity while completing the task. A temporal response function (TRF) was then trained by that EEG data and two regressors of the continuous speech stimulus. These two regressors were the speech envelope and gammatone spectrogram of the narrative. These backward TRF models were then used to predict the speech envelope and gammatone spectrogram based on the EEG data. A Pearson correlation coefficient was then calculated to evaluate how well correlated the predicted and actual values were.

The participants heard the same story twice, once as the story they were attending to and another time as the story, they ignored while attending to a different narrative. The blue trends show the TRF models for the attended conditions while the red trends show the models for the unattended conditions. Each of these show how well the brain is able to entrain on the different features of continuous speech.

Figure 2 is a boxplot of the correlation coefficients for the different backwards TRF models predicting for speech envelope and the gammatone spectrogram of the continuous speech narrative across the attended and unattended conditions. A two-way ANOVA and Tukey multiple comparison post-hoc analysis were used to test for the significant differences of TRF R-values across conditions and regressors. For the speech TRF envelope model, the attended condition (M= 0.1227 ± 0.05234) was significantly greater (F(1, 96) = 73.171, p< 0.0001) than the unattended conditions (M= 0.05087 ± 0.0242). The same pattern was also seen for the gammatone spectrogram model where the attended condition (M= 0.07712 ± 0.03217) was significantly greater (F(1, 96) = 73.171, p< 0.001) than the unattended condition (M= 0.03339 ± 0.01434). There was also a main effect across conditions where the envelope model R-values were significantly greater than the gammatone models (F(1, 96) = 21.789). More specifically, there were significant group difference between the attended envelope model and both of the conditions of the gammatone model (F(1, 96) = 4.327, p_attended_ < 0.001, p_unattended_ < 0.0001). The unattended envelope model was significantly smaller than the attended gammatone model (F(1, 96) = 4.327, p= 0.03549) but was not significantly different from the unattended gammatone condition (F(1, 96) = 4.327, p= 0.2658).

**Figure 2.**
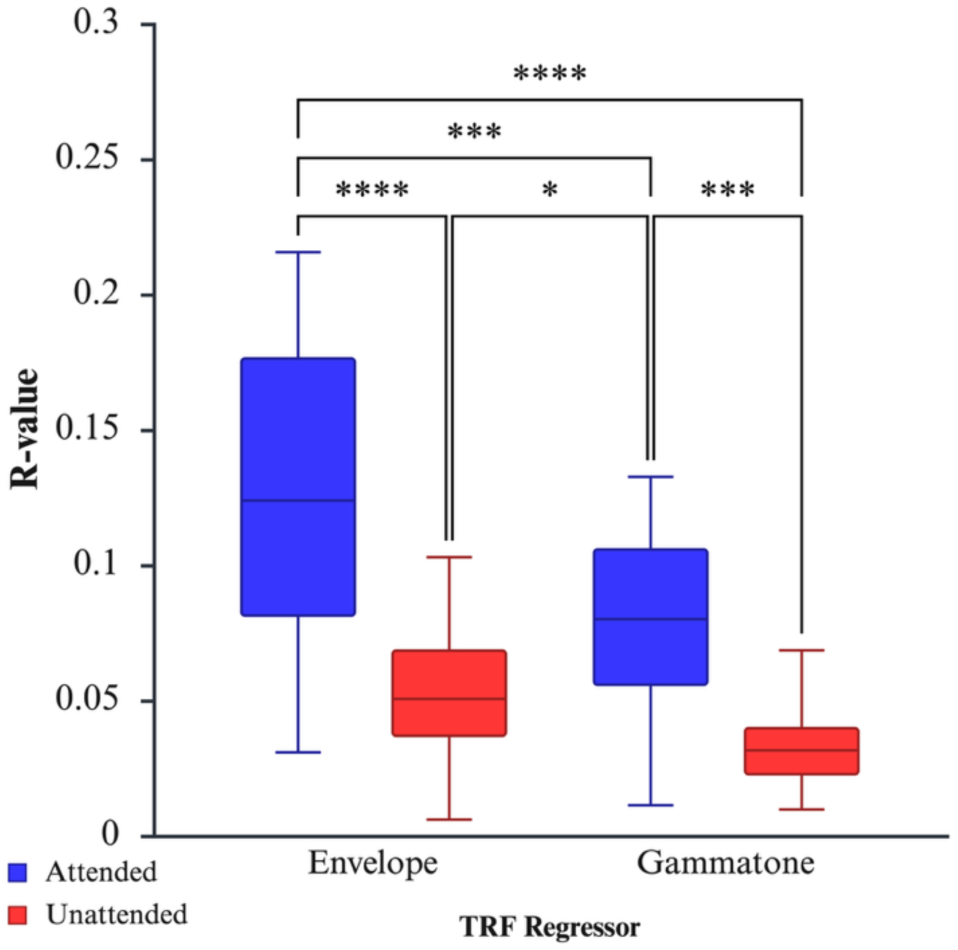
Box plots of temporal response function (TRF) cross validation correlation coefficient (R-values) across different task condition and TRF model regressors. ANOVA and Tukey multiple comparison post-hox analysis were used to test for group differences. * p < 0.05, *** p < 0.001, **** p < 0.0001

### Correlation of Neural Entrainment and Task Performance

Multiple regression models were used to evaluate the relationship between task performance and how well the brain entrained to the speech envelope of either an attended or unattended stimulus. Task performance was predicted using four predictor variables: TRF correlation coefficients from the attended condition, TRF correlation coefficients from the unattended condition, the subjects’ age, and the subjects’ pure tone audiometry (PTA) threshold averages. The participants’ ages and PTAs were used to test and control if those factors influenced the models.

### Entrainment of Speech Envelope and Task Performance Models

Hit accuracy was significantly correlated with the neural entrainment of the speech envelope for both the attended and unattended conditions. The attended conditions had a positive relationship where the greater R-value was correlated with better performance on the task. The inverse pattern was observed for the unattended condition where greater R-value corresponded with worse task performance. The participants’ age and PTA values were not significantly correlated with task performance (Figure 3.a, Table 1.a). Hit reaction time showed a similar pattern to hit accuracy. Increase attended R-values were significantly correlated with faster reaction times while there was a nonsignificant trend between slower reaction times and greater unattended R-values (Figure 3.b, Table 1.b). Neither age or PTA were correlated with hit RT. Narrative comprehension accuracy had a significant relationship with the attended TRF R-value but no relationship with the unattended TRF, participant age, or PTA values (Figure 3.c, Table 1.c).

**Figure 3.**
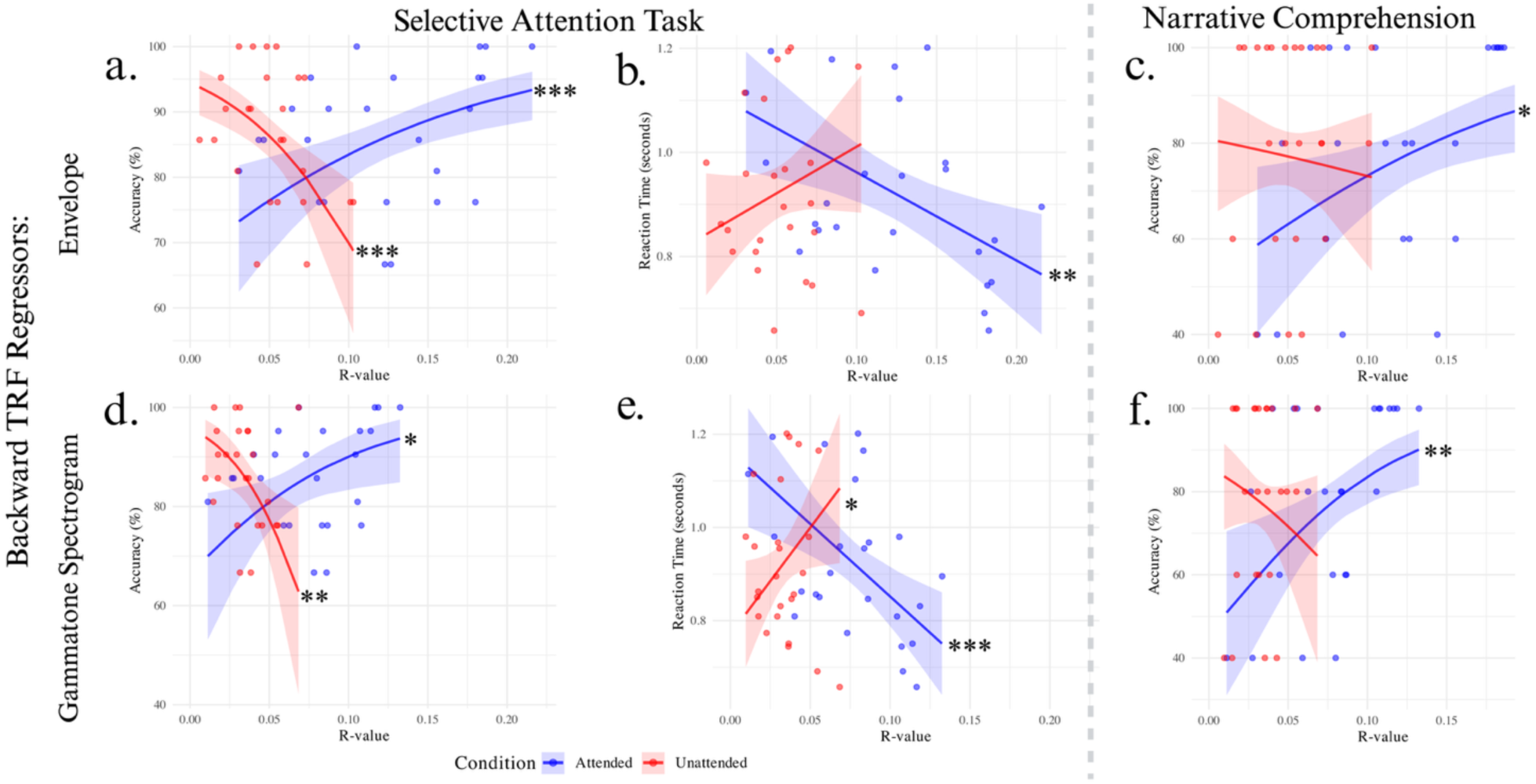
Scatter plots of TRF cross validation correlation coefficients (R-value) by speech-in-noise task performance. The top row (a-c) is the relationship between task performance and R-values from backwards TRF models predicting the speech envelope and the second row (d-f) is for the models predicting for the gammatone spectrogram. Each column is for the different task measures for a & d) hit accuracy on a spatial selective auditory attention task, b & e) hit reaction time on a spatial selective auditory attention task, and c & f) accuracy on a narrative comprehension task. Predictive modeling was performed using multiple regression models. * p < 0.05, ** p < 0.01, *** p < 0.001,

**Table 1.** Multiple regression models where task performance is predicted by the correlation coefficients of the backwards TRF model predicting for the speech envelope of the narrative stimulus. The three sub Tables are the models for each measure of task performance and their respective model type: a. color word hit accuracy (beta regression), b. color word hit reaction time (RT) (linear regression), b. narrative comprehension accuracy (beta regression).

| a. Hit Accuracy | Estimate | Std. Error | z value | p |
| --- | --- | --- | --- | --- |
| (Intercept) | 1.82160 | 0.12721 | 14.319 | < 0.001 |
| Attended TRF R-Value | 0.46332 | 0.13840 | 3.348 | < 0.001 |
| Unattended TRF R-Value | -0.48018 | 0.12806 | -3.750 | < 0.001 |
| Age | 0.10003 | 0.12848 | 0.779 | 0.436235 |
| PTA | 0.08622 | 0.11681 | 0.738 | 0.460420 |
| **b. Hit RT** | **Estimate** | **Std. Error** | **z value** | **p** |
| (Intercept) | 0.92316 | 0.02517 | 36.68 | < **0.001** |
| Attended TRF R-Value | -0.08875 | 0.03010 | -2.95 | < **0.01** |
| Unattended TRF R-Value | 0.04330 | 0.02913 | 1.49 | 0.13717 |
| Age | -0.04280 | 0.02656 | -1.61 | 0.10705 |
| PTA | -0.02300 | 0.02755 | -0.83 | 0.40381 |
| **c. Comp Accuracy** | **Estimate** | **Std. Error** | **z value** | **p** |
| (Intercept) | 1.2178 | 0.1636 | 7.444 | < **0.0001** |
| Attended TRF R-Value | 0.4916 | 0.1956 | 2.513 | **0.012** |
| Unattended TRF R-Value | -0.1060 | 0.1894 | -0.560 | 0.575 |
| Age | 0.1103 | 0.1726 | 0.639 | 0.523 |
| PTA | 0.1597 | 0.1791 | 0.892 | 0.373 |

### Entrainment of Gammatone Spectrogram and Task Performance Models

The trends observed across the three task measurements and the backwards TRF models of the speech envelope were the same as the trends observed for the models predicting the gammatone spectrogram. While the trends were the same, there were noTable differences in the strength of the trends across task measures.

Hit accuracy still has a significant positive relationship between the attended and a significant negative relationship with the unattended TRF correlation coefficients, but they are not as strong as the speech envelope models (Figure 3.d, Table 2.a). The inverse is true for the hit RT measure. The significant relationship between increased R-values and faster RT is now more significant. In addition, the trend between the greater unattended R-values and slower RT was not significant for the speech envelope TRF models but was significant for the gammatone spectrogram TRF models (Figure 3.e, Table 2.b). This pattern was also seen for the narrative comprehension question performance where only the attended TRF R-values were even more significant and there was still no trend for the unattended TRF R-values and task performance (Figure 3.f, Table 2.c).

**Table 2.**
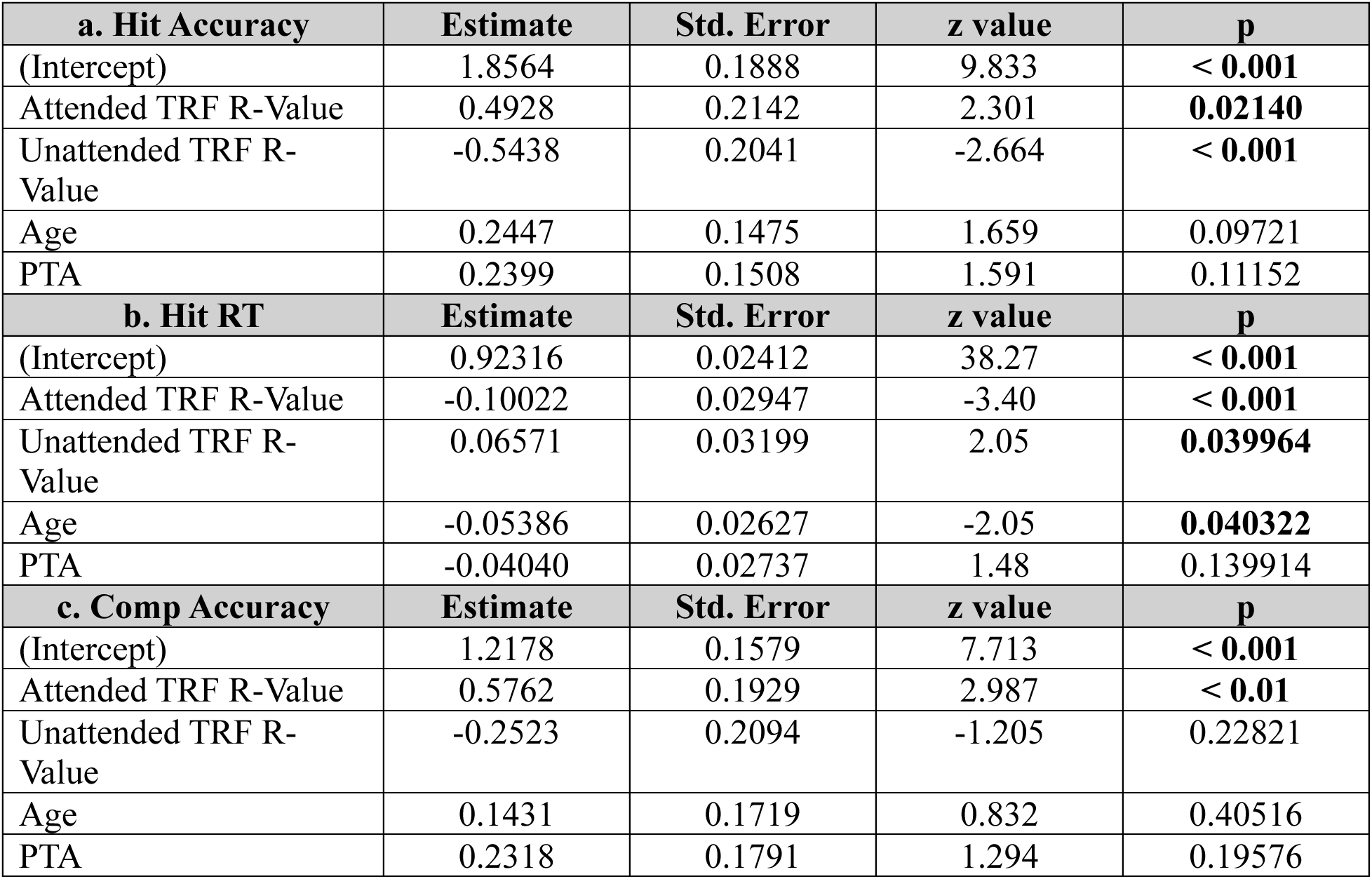
Multiple regression models where task performance is predicted by the correlation coefficients of the backwards TRF model predicting for the gammatone spectrogram of the narrative stimulus. The three sub Tables are the models for each measure of task performance and their respective model type: a. color word hit accuracy (beta regression), b. color word hit reaction time (RT) (linear regression), b. narrative comprehension accuracy (beta regression).

Age and PTA were not significant for any of the gammatone spectrogram backwards TRF models, with the exception of hit reaction time. This model showed that age had a negative relationship with hit RT. This means older participants had faster reaction times than younger participants. To understand if this could be due to a possible interaction between the TRF R-values and age, mixed effects modeling was used. The main effects of the R-values remained significant but there was no significant interaction between the R-values and age. This model also no longer showed a significant relationship between age and RT (Table 3).

**Table 3.** Linear mixed effects modeling to test for intractions of age for color word hit reaction time predicted by backwards TRF gammatone model.

| **Hit RT** | **Estimate** | **Std. Error** | **z value** | **p** |
| --- | --- | --- | --- | --- |
| (Intercept) | 0.93439 | 0.02419 | 38.63 | < **0.001** |
| Attended TRF R-Value | -0.09657 | 0.02855 | -3.38 | < **0.001** |
| Unattended TRF R-Value | 0.07601 | 0.03189 | 2.38 | **0.017135** |
| Age | -0.02471 | 0.03239 | -0.76 | 0.445574 |
| PTA | -0.03837 | 0.02621 | 1.46 | 0.143296 |
| Attended TRF:Age | 0.02864 | 0.03059 | 0.94 | 0.349113 |
| Unattended TRF:Age | -0.04135 | 0.02687 | -1.54 | 0.123786 |

**Table 4.** Multiple regression models where task performance is predicted by the difference of correlation coefficients across conditions (attended -unattended) of the backwards TRF model predicting for the speech envelope of the narrative stimulus. The three sub Tables are the models for each measure of task performance and their respective model type: a. color word hit accuracy (beta regression), b. color word hit reaction time (RT) (linear regression), b. narrative comprehension accuracy (beta regression).

| **a. Hit Accuracy** | **Estimate** | **Std. Error** | **z value** | **p** |
| --- | --- | --- | --- | --- |
| (Intercept) | 1.79808 | 0.13719 | 13.107 | < **0.001** |
| TRF R-Value Difference | 0.41911 | 0.13009 | 3.222 | < **0.01** |
| Age | 0.12670 | 0.14337 | 0.884 | 0.37682 |
| PTA | 0.03208 | 0.12331 | 0.260 | 0.79472 |
| **b. Hit RT** | **Estimate** | **Std. Error** | **z value** | **p** |
| (Intercept) | 0.92316 | 0.02517 | 36.67 | < **0.001** |
| TRF R-Value Difference | -0.07841 | 0.02655 | -2.95 | < **0.01** |
| Age | -0.04298 | 0.02649 | -1.62 | 0.10474 |
| PTA | -0.02257 | 0.02712 | -0.83 | 0.40532 |
| **c. Comp Accuracy** | **Estimate** | **Std. Error** | **z value** | **p** |
| (Intercept) | 1.02616 | 0.15277 | 6.717 | < **0.001** |
| TRF R-Value Difference | 0.33335 | 0.15785 | 2.112 | **0.0347** |
| Age | 0.08764 | 0.16282 | 0.538 | 0.5904 |
| PTA | 0.12655 | 0.14689 | 0.862 | 0.3889 |

### Relationship of TRF Correlation Coefficient Differences and Task Performance

Both hit accuracy and hit RT showed an inverse relationship across the attended and unattended conditions. Better performance on the task was positively correlated with the attended TRF R-values and negatively correlated with the unattended R-values. To further investigate this pattern, the difference of the R-values (attended – unattended) were used to predict task performance.

Increased task performance was correlated with a greater difference between TRF R-values for all three task performance measurements. The difference of TRF correlation coefficients of the speech envelope models had a positive relationship between hit accuracy (Figure 4.a, Table 5.a), a negative relationship with RT (aka faster response time) (Figure 3.b, Table 5.b), and a positive relationship with narrative comprehension accuracy (Figure 3.c, Table 5.c). The same trends with the gammatone spectrogram TRF models were also observed for hit accuracy (Figure 3.d, Table 6.a), hit reaction time (Figure 3.e, Table 6.b), and narrative comprehension accuracy (Figure 3.f, Table 6.c). These models showed the same degree of significant differences as previous models where the speech envelope model best predicted for hit accuracy while the gammatone spectrogram best predicted for hit RT and narrative comprehension performance.

**Figure 4.**
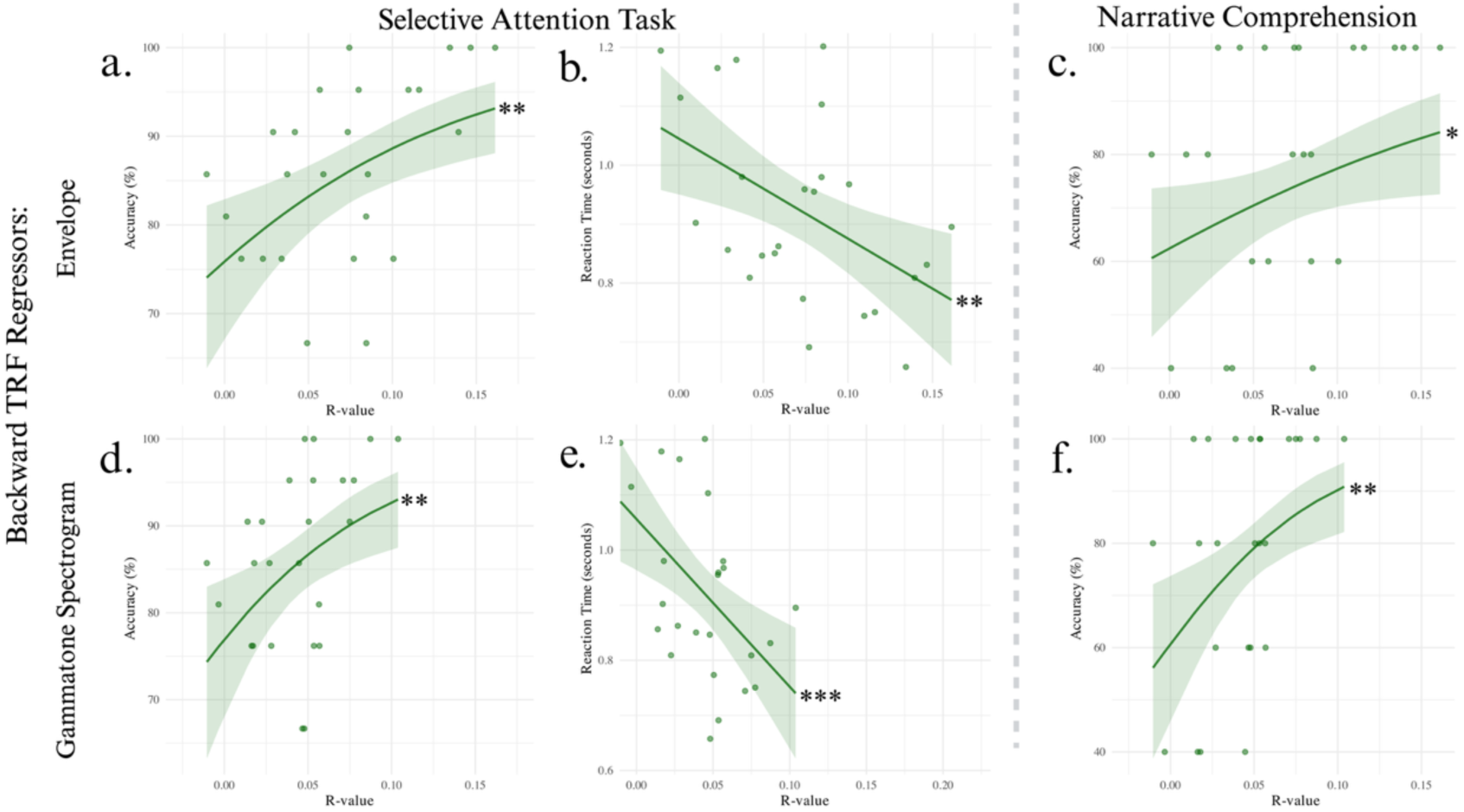
Scatter plots of the differences (attended – unattended conditons) of TRF cross validation correlation coefficients (R-value) by speech-in-noise task performance. The top row (a-c) is the relationship between task performance and R-values from backwards TRF models predicting the speech envelope and the second row (d-f) is for the models predicting for the gammatone spectrogram. Each column is for the different task measures for a & d) hit accuracy on a spatial selective auditory attention task, b & e) hit reaction time on a spatial selective auditory attention task, and c & f) accuracy on a narrative comprehension task. Predictive modeling was performed using multiple regression models. * p < 0.05, ** p < 0.01, *** p < 0.001,

**Table 5.** Multiple regression models where task performance is predicted by the difference of correlation coefficients across conditions (attended -unattended) of the backwards TRF model predicting for the gammatone spectrogram of the narrative stimulus. The three sub Tables are the models for each measure of task performance and their respective model type: a. color word hit accuracy (beta regression), b. color word hit reaction time (RT) (linear regression), b. narrative comprehension accuracy (beta regression).

| **a. Hit Accuracy** | **Estimate** | **Std. Error** | **z value** | **p** |
| --- | --- | --- | --- | --- |
| (Intercept) | 1.78823 | 0.14019 | 12.756 | < **0.001** |
| TRF R-Value Difference | 0.36865 | 0.12725 | 2.897 | < **0.01** |
| Age | 0.15998 | 0.14534 | 1.101 | 0.27099 |
| PTA | 0.08569 | 0.12773 | 0.671 | 0.50231 |
| **b. Hit RT** | **Estimate** | **Std. Error** | **z value** | **p** |
| (Intercept) | 0.92316 | 0.02440 | 37.84 | < **0.001** |
| TRF R-Value Difference | -0.08391 | 0.02541 | -3.30 | < **0.001** |
| Age | -0.04959 | 0.02594 | -1.91 | 0.055955 |
| PTA | -0.03262 | 0.02565 | -1.27 | 0.203416 |
| **c. Comp Accuracy** | **Estimate** | **Std. Error** | **z value** | **p** |
| (Intercept) | 1.2178 | 0.1580 | 7.707 | < **0.001** |
| TRF R-Value Difference | 0.4943 | 0.1644 | 3.006 | < **0.01** |
| Age | 0.1440 | 0.1680 | 0.858 | 0.39112 |
| PTA | 0.2334 | 0.1660 | 1.406 | 0.15970 |

## Discussion

The aim of this investigation was to see if metrics of neural entrainment using TRF are related to the performance on a speech-in-noise spatial selective attention task. EEG data was collected while participants completed an ecologically relevant spatial auditory attention task where they attended to a narrative of continuous speech. This was performed on a collection of young adults with clinically “normal” hearing. This means we were able to better control for age and hearing ability when investigating for the variance of perceptual ability in a cocktail party environment.

### Task Performance

Participants performed well on all three measures collected from our spatial auditory attention task and narrative comprehension task. Participants’ hit accuracy averaged around 86% and varied around 10% in either direction. This illustrates that the task was not too easy, but it was also challenging enough to show a spread of SIN perceptual performance. The same is also true for the hit RT measure that most participants ranged around 1 second. These are both measures of the participants’ ability to selectively attend to the target while ignoring the masker narrative and we were able to demonstrate the unexplained varied nature of SIN perception in a young and healthy cohort.

After completing the spatial auditory attention task, participants then answered narrative comprehension questions about the events of the attended narrative. Participants performed well with many of them achieving perfect scores. This number of high-performers could be in indicator of either the questions being too easy or the need of more questions to prevent possible ceiling effects. Our analysis still observed significant trends for the measure because the wide range of performances across the whole cohort and the uses of non-linear beta regression modeling to compensate for possible ceiling effects.

### TRF Correlation Coefficients as a Measure of Neural Entrainment

The correlation between a predicted variable using TRF models trained on neural data and the actual signal has been well documented to be related to attention.^9^ This has been interpreted as different aspects of an auditory signal (e.g., the envelope and spectral energy of a sound wave) having a linear relationship with different neural processes that entrain onto those specific features. While these correlation values are extremely small (R-values ranging from 0.05 to .3), they are still significant and modulated by high-level cognitive processes.^8^ A participant’s TRF prediction correlation coefficient will be significantly greater while they attend to a stimulus then their R-values while ignoring an identical stimulus.

TRF investigations are most commonly performed with forward models where regressors from a stimulus are used to predict the EEG waveform of every electrode across the scalp.^15^ We decided to use backwards TRF models for this investigation instead. We predicted the speech envelope and gammatone spectrogram of our auditory stimulus using the participants’ EEG waveforms from a dense array of 64-electrodes. The motivation behind this decision was to find a concise measure of neural entrainment that is representative of the neural activity across the whole scalp.

We were able to reproduce the attentional findings from other TRF analyses, where the TRF R-values for our attended conditions were significantly greater than the unattended conditions. The R-values were also more variable for the attended conditions than the unattended conditions. This was true for both the speech envelope and gammatone spectrogram models.

In addition to a main effect across the conditions for the two regressor models, we observed a significant difference across the two regressor models themselves. There was a main effect across the regressor models where the envelope R-values were significantly greater than the gammatone spectrogram models. Surface level assessment of the magnitude of the different R-values would imply that speech envelope is a better regressor when predicting for auditory attention, but this is not observed when correlating TRF R-values with task performance.

### The Relationship Between Neural Entrainment and Speech-in-Noise Perception

Differences in R-values across conditions illustrates that the brain will better entrain onto a stimulus when directing its cognitive resources to an auditory target, but the same degree of neural entrainment is not observed for an unattended stimulus. While the correlation coefficients are significantly smaller, the unattended stimulus still retains a small degree of neural entrainment. This aligns with ERP analysis where an auditory stimulus will evoke an obligatory response for both attended and unattended stimuli.^16^ While they both elicit a response, the ERP morphology will differ across conditions. The ERPs to the attended stimulus will have a greater amplitude and smaller latencies than the ignored stimulus. We believe the same principle is being observed with TRF where the brain will entrain to both signals, but will modulate the degree of entrainment based on the participant’s attention.

It has been well documented that the brain entrains to both attended and unattended stimuli,^8^ but we aim to investigate if the degree of neural entrainment is correlated with one’s ability to perceive speech in noisy environments. We performed this analysis on not just the attended trials but also the unattended trials. We posit that the successful listeners in a cocktail party environment would simultaneously have increased neural entrainment on the target narrative while having a more greatly suppressed entrainment on the distractor stimulus. The logic being that the person’s brain would have a robust neural representation of the auditory target while having a weaker and suppressed representation of the distractors. These distinct neural representations would enable a participant to more easily identify the target color words in the narrative and can better comprehend the information from the narrative, which would increase their performance on the three behavioral tasks in this study.

This model would also imply that poor performers in noisy environments will have TRF R-values across conditions that are closer to each other. This would result in a noisier and possibly overlapping representation of the two competing narratives. The lack of ability to parse the two narratives from each other could result in diminished SIN performance because it is more difficult to extract the information from the target narrative if there is an equally represented distractor narrative competing for the same cognitive resources. This would affect behavior by making it more difficult to identify and respond to the color words in the target narrative and would have a diminished ability to remember details from the narrative which are needed to answer the comprehension questions.

Our hypothesized model was observed in both behavioral metrics of the spatial selective attention task. Color word hit accuracy had had a significant positive relationship with the attended TRF R-values and negative relationship with unattended TRF R-values. This was observed for both regressor models, but with a stronger correlation for the speech envelope model. Color word hit RT was also negatively correlated with attended conditions while having a positive relationship with the unattended condition. The relationship between RTs and attended R-values were significant for both models, but unattended R-values were significantly correlated with the gammatone spectrogram models and had a non-significant trend for the speech envelope models. This means the participants with the highest accuracy and fastest RTs are the ones with the highest attended R-values and lowest unattended R-values while the worst performers have the smallest attended and largest unattended TRF R-values.

This model was not completely observed for participants’ performance on the narrative comprehension questions. Rather than both R-values being correlated, only the attended R-values were related to narrative comprehension accuracy.

These finding support our model that neural entrainment to the attended narrative is related to how well a person is able to perceive a target. In addition, that degree of suppression of the unattended stimulus is only significant during the selective attention task. The suppression of neural entrainment during the unattended conditions is not significantly related to narrative comprehension. We posit that is because the different tasks require different neural mechanisms. While actively attending to the target stimuli and ignoring the masker, it is equally important to robustly represent the target and suppress a response to the unattended stimulus. This means that participants who have a strong differentiation across conditions are able to be more accurate and faster in their responses to the stimulus. This differentiation across streams of information is not as significant when encoding the information that would be needed to recall after the task to answer the narrative comprehension questions. While there was no relationship across comprehension performance and the unattended TRF R-values, there was a strong correlation with the attended R-values. Participants who were able to better entrain on the target narrative were better able to answer the comprehension questions. Meaning the overall strength of entrainment to a stimulus is strongly associated with increased narrative comprehension.

Correlation coefficients across backwards TRF regressor models were significantly different from each other. The R-values of the speech envelope were significantly larger than the R-values for the gammatone spectrum model. Despite the significantly lower R-values, gammatone spectrogram models were still correlated with task performance and even had greater correlations with two performance metrics than the larger speech envelope R-values. RT and narrative comprehension were more significantly correlated with the gammatone. This increased correlation could be due to the nature of the information being encoded with each regressor, such as greater entrainment of the envelope could make it easier for a participant to glimpse the target narrative in the gaps or a greater representation of the gammatone spectrogram allows the participants to better perceive the formants of a word so they can better process the narrative meaning of the words in the narrative. Further investigation of how different regressors relate to behavior is necessary.

### Differences of Neural Entrainment as a Biomarker of Attention

The difference of R-values across conditions was calculated to isolate the effects of attention on neural entrainment. The stimulus across conditions was identical and the only difference was which narrative the participant was directed to attend to. This means that differences in R-values is primarily due to how the participant directed their attention. This metric was significantly correlated with all three task performance metrics for both the speech envelope and gammatone spectrogram regressor models. Showing that greater degree of neural entrainment across the conditions is correlated with greater ability to succeed in a cocktail party environment.

Showing the differences across R-values is a robust measure for predicting task performance. This measurement could be used as a biomarker for future investigations and hopefully clinical use.

### Limitations and Future Directions

It is possible that differences in neural entrainment could be one of the mechanisms responsible for the wide range of speech perception ability. Further investigation of these differences is necessary to help us create a more comprehensive understanding of the neural mechanisms underlining the cocktail party problem. Future studies should specifically evaluate if differences in neural entrainment could be used as a biomarker to identify those who might experience deficits in speech perception in a noisy environment.

This investigation was a part of a larger project to characterize the hearing ability of a large cohort of military veterans. Future analyses will include investigating the differences in TRF R-values in the cohort that is comprised of over one-hundred participants who greatly vary in their ages, clinical hearing ability, cognitive ability, and a wealth of other factors. The added stress on the neural mechanisms of speech perception due to a less salient auditory signal could more severely impact those with smaller TRF R-value differences, which could be a possible mechanism for poorly defined deficits like hidden hearing loss.

Furthermore, differences in neural entrainment might also be related to different auditory perception deficits associated with disorders like ADHD and auditory processing disorder (APD). Further studies with neurodivergent cohorts could help better understand the relationship of neural entrainment and auditory attention deficits.

Current findings were only reported on how backwards TRF models can be used as a biomarker to predict a person’s ability to succeed in a cocktail party environment. Future analyses should investigate using forward TRF models relationship with task performance. Specifically investigating if different regions of the scalp are correlated with different performance metrics. The TRF filters should also be evaluated to see if their morphology aligns with these findings.

Finally, the narrative stimuli were presented with Cheech rather than normal speech. This method can effectively investigate the neural activity from cochlea to cortex by evoking auditory brainstem responses, middle latency responses, and late latency responses in ERP analyses. The chirped embedded in the Cheeched stimulus could be a regressor to evaluate the brainstem’s role in auditory attention and neural entrainment.

## Acknowledgements

Funding for this research was provided by the Department of Defense (DoD) and the Child Family Fund for the Center for Mind & Brain. The authors thank the audiology and clinical research team of the University of California, Davis Health for conducting initial participant hearing evaluations including, Dr. Robert Ivory, Au.D., Dr. Mackenzie Quinn, Au.D., Dr. Rachel Krager, Au.D., Dr. Steven Zurawski, Au.D., Dr. Austin Childers, Au.D., Dr. Kimberly Smith, Au.D., Randev Sandhu, and Angela Beliveau. We also thank Cathleen Chan and Jillian McKie for their contributions to data collection, and Elyse Ehlert and Tiana Smith for their work in recruiting our participants. Additionally, we are grateful for valuable discussions with Sophie Burstein, Alicia Dye, Reina Itakura, Zacharay McNaughton, Ferdous Rahimi, Tyler Statema, Sana Shehabi, and Audrey Vargas. Most importantly, we extend our deepest gratitude to all of the participants whose time and effort made this research possible. This work is dedicated to the memory of our team member, lab mate, and friend, Karim Abou Najm.

## Author Contributions

D.C.C, K.M, H.B, D.S, and L.M.M designed research; B.B, and S.D performed research; D.C.C, K.M, B.B, Z.B, C.L, A.G, R.W and L.M.M contributed analytic tools; B.B, Z.B, C.L, A.G, and D.C.C analyzed data; B.B, Z.B, D.C.C, K.M, R.W, S.D, and L.M.M wrote and edited the paper.

